# Kaempferide induces apoptosis in cervical cancer by attenuating the HPV oncoproteins, E6 and E7

**DOI:** 10.1101/2023.05.19.541414

**Authors:** Lekshmi R Nath, Vijai V Alex, Sreekumar U Aiswarya, Tennyson P Rayginia, Nair Hariprasad Haritha, Chenicheri K. Keerthana, Arun Kumar T. Thulasidasan, Mundanattu Swetha, Ravi Shankar Lankalapalli, Ruby John Anto

## Abstract

**Graphical Abstract.**
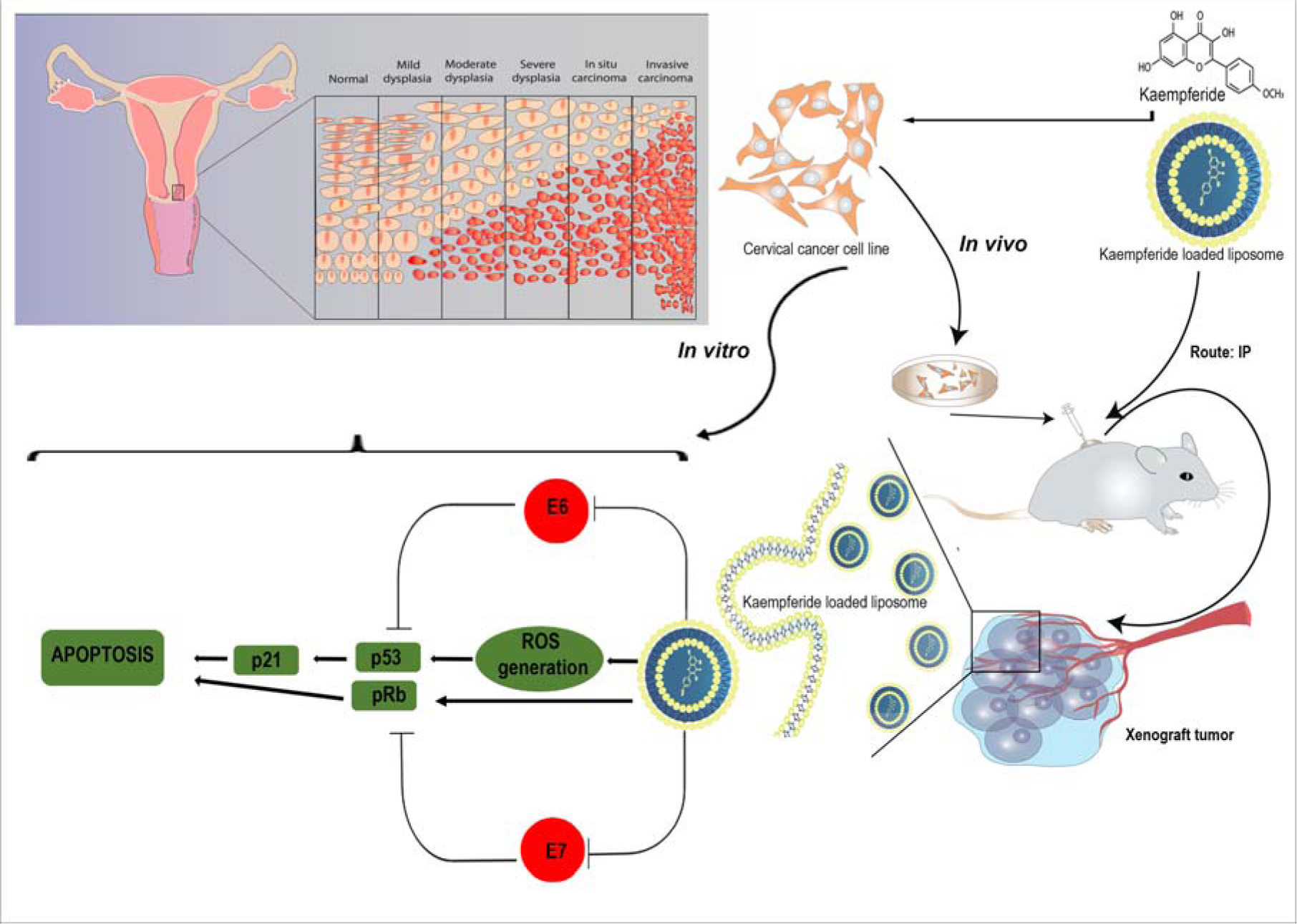
A schematic diagram depicting the anti-cancer potential of kaempferide against cervical cancer and its probable mechanism of action.

Despite the advancement in HPV prevention strategies, cervical cancer is a leading cause of cancer death in women worldwide. The anticancer potential of kaempferide, which was derived from *Chromolaena odorata*, was previously revealed in our in-vitro study. The current investigation aims to confirm the therapeutic efficacy of the molecule against cervical cancer, *in-vitro* and *in-vivo*. In NOD-SCID mice with HeLa Luc+ xenografts, kaempferide significantly increases ROS production resulting in a drastic decrease in the luc activity and growth of cervical tumours. It also exhibits down-regulation of oncoproteins E6, E7 and MDM2 and concurrent up-regulation of p53, p21 and pRb. The degradation of phospho-pRb, along with a strong expression of cleaved PARP, TUNEL positivity and a significant decrease in PCNA expression, confirms apoptotic mode of cell death in the treated tissues. Taken together, our study reveals the antioncogenic potential of kaempferide, which regulates oncoproteins and tumor suppressors accordingly. This is the first report depicting kaempferide as an inhibitor of HPV oncoproteins and hence as a candidate drug molecule against cervical cancer.

## 1. Introduction

Cervical cancer is the fourth most common cancer in women worldwide. Though the advancement in early detection through screening strategies and vaccination programs has reduced the incidence and mortality of cervical cancer in developed countries, it is still a significant concern in developing countries as these programs are not much accessible to the victims due to the high cost. Moreover, the available vaccines are not useful to the infected women, women above 27 and women, who are already sexually active before the vaccination.

The causative role of persistent infection by High-risk human papilloma viruses (HR-HPVs) is a well-known factor in cervical cancer progression [1]. The predominant role of the HPV proteins, E6 and E7 and their contribution in altering the signaling mechanisms, which lead to cellular transformation and carcinogenesis is well understood. Evasion of cell cycle checkpoints is a crucial process in tumorigenesis. HPV is known to target cellular factors that regulate cell cycle checkpoints. P53 and pRb are well known cellular factors that deregulate cell cycle upon stress or other tumorigenic phenotypes. HPV viral proteins E6 and E7 target and degrade these two tumor suppressors, aiding cell cycle checkpoint evasion [2], enabling the transformation of healthy cells to a proliferative state. Reports claim that down-regulating the expression of these viral proteins using different molecular techniques can reverse this proliferative nature and induce apoptosis [3, 4].

Cervical cancer, if detected at an early stage, is considered to be generally curable. However, treatment regimens against metastatic or recurrent carcinoma are usually having abysmal prognosis and hence cause serious side effects. For advanced stages, the first choice of treatment is still chemotherapy. Combination of chemotherapy, surgery and radiation significantly improve the survival of the patient. But, the development of chemoresistance and radioresistance combined with adverse side effects pose serious negative effects on the quality of life and treatment outcome for the patient. These facts are pointing towards the importance of developing novel therapeutic strategies against cervical cancer [5, 6].

A large group of plant-derived polyphenols, especially flavonoids, contain a broad spectrum of secondary metabolites such as chalcones, flavones, flavanols, flavanones and isoflavones, which have been reported to be cytotoxic to many cervical cancer cell lines. Some are known to induce apoptosis in cervical cancer cell lines [7–9] and some others work via generation of ROS [10–12] and regulating the cell cycle [13]. Several studies have also reported the potential of flavonoids to suppress the expression of E6 and E7 [14–16].

*Chromolaena odorata*, commonly known as Siam weed, is a widely popular plant in traditional medicine. Our lab isolated four flavonoids from the DCM extract of the plant, among which, kaempferide was found to be the most potent compound, which exhibited maximum efficacy against cervical cancer, while being pharmacologically safe towards normal cells and healthy animals [17]. The present study was conducted to evaluate the mechanism of action of kaempferide and confirm the efficacy of the molecule against cervical cancer *in vivo*, using a cervical cancer xenograft model.

## 2. Materials and Methods

### 2.1 Reagents and antibodies

Cell culture reagents such as Dulbecco’s Modified Eagle Medium (DMEM) (GIBCO, 12800-017) and streptomycin sulfate (GIBCO, 11860-038) were obtained from Invitrogen Corporation (Grand Island, USA). Poly Excel HRP/DAB detection system universal kit (PathnSitu Biotechnologies Pvt. Ltd, India, OSH001) was used for immunohistochemistry experiments. MTT reagent was purchased from TCI Chemicals (India) Pvt. Ltd (D0801) and Amersham ECL Plus™ Western blotting reagents (PRPN 2132) were purchased from GE Healthcare Life Sciences (Piscataway, USA). Antibodies against, p-P53 (9281S), PARP (9532S), β-actin (12620S) were obtained from Cell Signalling Technologies (Beverly, MA, USA) and the antibody against PCNA (sc25280) were purchased from Santa Cruz Biotechnology (Santa Cruz, CA, USA). and ECL reagent (Pierce™ ECL western blotting substrate 32109) were purchased from ThermoFisher Scientific (Waltham, Massachusetts, United States) Cellular ROS kit (ab113851) were purchased from Abcam (Cambridge, United Kingdom). DeadEnd™ Colorimetric TUNEL System from Promega (G7132) was procured from Addgene (Cambridge, MA, USA). Anti-rabbit and anti-mouse antibodies were obtained from Sigma Chemicals (St. Louis, MO, USA). All other chemicals and an antibody against Vinculin (V9131) were purchased from Sigma Chemicals (St. Louis, MO, USA) unless otherwise mentioned.

### 2.2 Cell lines

HeLa cells were procured from NCCS, Pune. Cell lines were maintained in DMEM supplemented with FBS (10%) and incubated in 5% CO_2_ at 37°C. Mycoplasma tests were performed on parent cell lines every 6 months. Cell lines passage between 3-6 times post-revival, were used for all experiments.

### 2.3 MTT assay

Two thousand cells were seeded in a 96-well plate and incubated overnight followed by incubation with different concentrations of the compounds for 72 hours. MTT [3-(4,5-dimethylthiazol-2-yl)-2,5-diphenyl tetrazolium bromide] assay was used to analyze the viability of cells to various concentrations of kaempferide as described in earlier [17].

### 2.4 Clonogenic assay

For assessing the colony-forming potential of HeLa cells, 500 cells were seeded/well in a 6-well plate, incubated overnight and were exposed to different concentrations of the compounds for 72 hours. The cells were then allowed to proliferate in fresh media for 72 hours before fixation and staining with crystal violet.

### 2.5 Wound healing assay

Confluent cells were cultured in DMEM containing 10% FBS for 24 h and then wounded with a linear scratch by a sterile pipette tip. Cell debris was gently removed by washing the cells with 1X PBS. To observe the effect of compound on migration, the cells were treated with various concentrations of kaempferide. Images were captured at different time intervals under a phase contrast microscope.

### 2.6 Transfection

Luciferase transfected HeLa cells were developed to track the cells *in vivo* using *in vivo* live imager. The cells were stably transfected with pKT2/mCa-RLuc-IRE vector (Renilla-Luc) using the Lipofectamine LTX and Plus Reagent Kit (Invitrogen, USA) according to the manufacturer’s protocol.

### 2.7 *In vivo* studies

A cervical cancer xenograft study was performed in female NOD-SCID mice following the approved guidelines of the Institute Animal Ethics Committee of Rajiv Gandhi Centre for Biotechnology, Thiruvananthapuram (IAEC/721/RUBY/2018). Luc+ HeLa cells (5X10^6^) were subcutaneously injected into the flank region of the mice. 2 weeks post the cell injection, the animals were grouped into control and treatment groups. Kaempferide being hydrophobic was dissolved in Cremophor vehicle (Cremophor EL/ethanol-1:1 ratio), and diluted with sterile PBS (1:1). The treatment group was injected intraperitoneally with 75 mg/kg of kaempferide, twice weekly for one month. The tumor size was measured weekly and the corresponding tumor volume was calculated as per the formula, (length x width2)/2. At the end of the experiment, the animals were euthanized and tumor tissues were collected. All tumor reduction studies using kaempferide were carried out according to prescribed guidelines, which were approved by the Institutional Animal Ethical Committee, RGCB.

### 2.8 Live *In Vivo* imaging

The representative animals from different groups were anesthetized by isoflurane inhalation and analysed for the development of tumor or the reduction in tumor growth and metastatic spread of the tumor cells to different organs using the live animal imager [The IVIS spectrum Non-invasive multimode small animal imager, Caliper Life Sciences, USA], after intraperitoneal injection of 0.275mM Renilla lucifersaw substrate [Viviren, Promega] dissolved in sterile PBS containing 0.1% BSA. Bioluminescence in the tumor region can be visualized in the live animals.

### 2.9 Histopathology

For histopathological examination, 5 μm sections cut from the formalin-fixed, paraffin-embedded tissues were deparaffinized, rehydrated, stained with hematoxylin and eosin and mounted with distyrene plasticizer and xylene (DPX) mountant [18].

### 2.10 Immunohistochemistry

Immunohistochemistry of specific proteins in the tissue sections was done using the Path in situ kit following the manufacturer’s instructions. Photomicrographs were captured using a Nikon Eclipse microscope equipped with Image-Pro Plus software.

### 2.11 TUNEL assay

TUNEL assay was performed to detect apoptosis in formalin fixed, paraffin-embedded xenograft tumor tissue sections using DeadEnd Colorimetric TUNEL System (Promega) following the manufacturer’s instructions.

### 2.12 Cellular ROS assay

The production of reactive oxygen species (ROS) within the cells in response to kaempferide were determined by staining the cells using H2DCF-DA according to the manufacturer’s protocol.

### 2.13 Statistical Analysis

All data were expressed as mean ± SD. Data were analyzed with Prism 6.0 (GraphPad Software). One-way ANOVA measured statistical significance between the conditions. **** P-values≤0.0001, ***P-values ≤0.001, **P-values ≤0.01 and *P ≤0.1 and ns ≥ 0.05

## 3. Results

### 3.1 Kaempferide exhibits anticancer potential against cervical cancer by down-regulating E6 and E7 oncoproteins, leading to p53 and pRb up-regulation

We previously reported for the first time, the cytotoxic potential of kaempferide, which was isolated in our lab, from the medicinal herb, *Chromolaena odorata*, against cervical cancer. The cytotoxic potential and purity of isolated kaempferide was re-confirmed by comparing with that of kaempferide purchased from Sigma, in the cervical cancer cell line, HeLa by MTT assay. Both samples of kaempferide showed a dose dependent cytotoxicity consistently, with an IC50 range around 15µM, **(Figure 1A)**, verifying our previous report [17]. Hence, we used the compound purchased from Sigma for all experiments. The clonogenic potential of HeLa cells was drastically inhibited by kaempferide as observed by more than 50% reduction in colony count at the IC50 concentration **(Figure 1B**). The compound also inhibited the migration of HeLa cells, at the IC50 concentration, as assessed by a significant reduction in wound closure in the scratch-wound healing assay compared to control wells, indicating the strong anti-migratory potential of kaempferide against cervical cancer **(Figure 1C)**.

**Figure 1:**
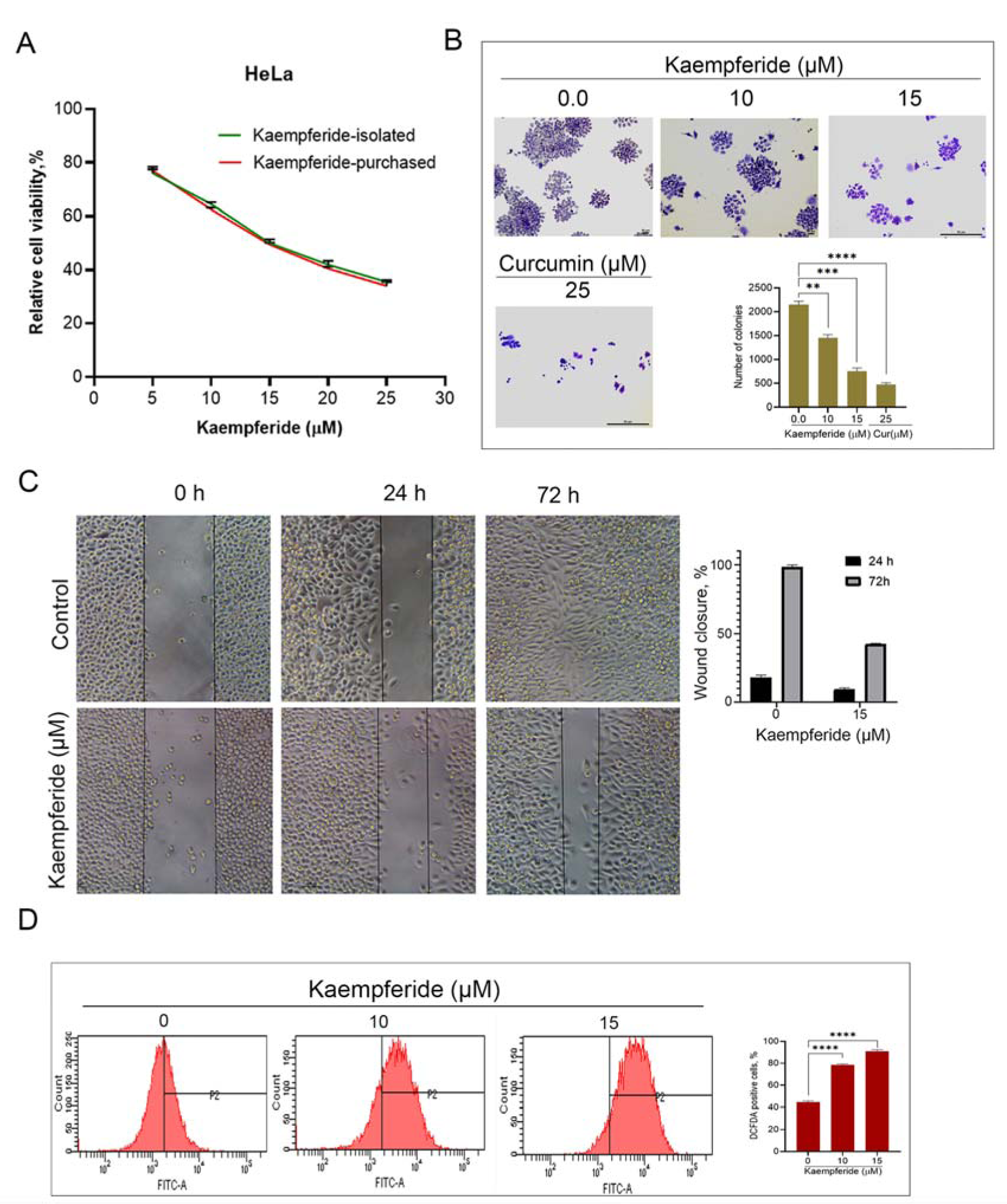
Kaempferide induces ROS production and inhibits the proliferation and migration of cervical cancer cells: **(A)** MTT assay to compare the cytotoxicity of isolated and commercial kaempferide in HeLa cells. **(B)** Clonogenic assay reveals the anti-clonogenic potential of kaempferide. **(C)** Wound healing assay showing the anti-migratory potential of kaempferide. **(D)** ROS production in response to kaempferide treatment in HeLa cells as detected by fluorescence microscopy. [Data are representative of three independent experiments (Mean±SEM) and P-values are calculated using one way ANOVA****P ≤0.0001,***P ≤0.001,**P ≤0.01,*P ≤0.1and ns ≥ 0.05].

Our previous study has demonstrated that kaempferide induces apoptotic mode of cell death in cervical cancer cells, independent of cell cycle [17] Several chemotherapeutic drugs have been reported to induce apoptosis in cancer cells by triggering DNA damage through induction of ROS production [19]. The ability of kaempferide to induce ROS production in HeLa cells was evaluated using an ROS-sensitive H2DCF-DA assay. Kaempferide treatment activated ROS production in HeLa cells in a dose-dependent manner, initiating the oxidation of H2DCF-DA to dichlorofluorescein (DCF), which further led to the generation of green fluorescence in the cells, the intensity of which was quantified by confocal microscopy demonstrating that kaempferide augments ROS production in cervical cancer cells, subsequently leading to DNA damage and intrinsic apoptosis **(Figure 1D)**.

Our next attempt was to elucidate the mechanism behind the high sensitivity of HeLa cells to kaempferide. HeLa cells over-express the HPV oncoproteins, E6 and E7, E6 being a key regulatory oncoprotein, which impairs apoptosis by targeting p53 degradation, while E7 inactivates the retinoblastoma susceptibility protein, pRb, in part by stimulating its degradation [20]. Interestingly, expression of E6 and E7 were strongly down-regulated by kaempferide as assessed by Western blot as well as gene expression studies **(Figure 2. A-C)**. Consequently, there was a strong up-regulation of p53 and its down-stream signaling molecule, p21, and concomitant down-regulation of MDM2, which inhibits p53 activities **(Figure 2D, E)**. This may be correlated to the strong ROS production, leading to p53-induced DNA damage. Overexpression of E7 causes proteasomal degradation of the tumor suppressor pRb, the phosphorylation of which exerts protection against apoptotic cell death in HPV-containing cervical cancer cells [21]. Subsequent to the down-regulation of E7 on kaempferide-treatment, we noted a strong up-regulation of pRb, and degradation of phospho-pRb, which may contribute to the apoptotic mode of cell death induced by the compound **(Figure 2F, G)**.

**Figure 2:**
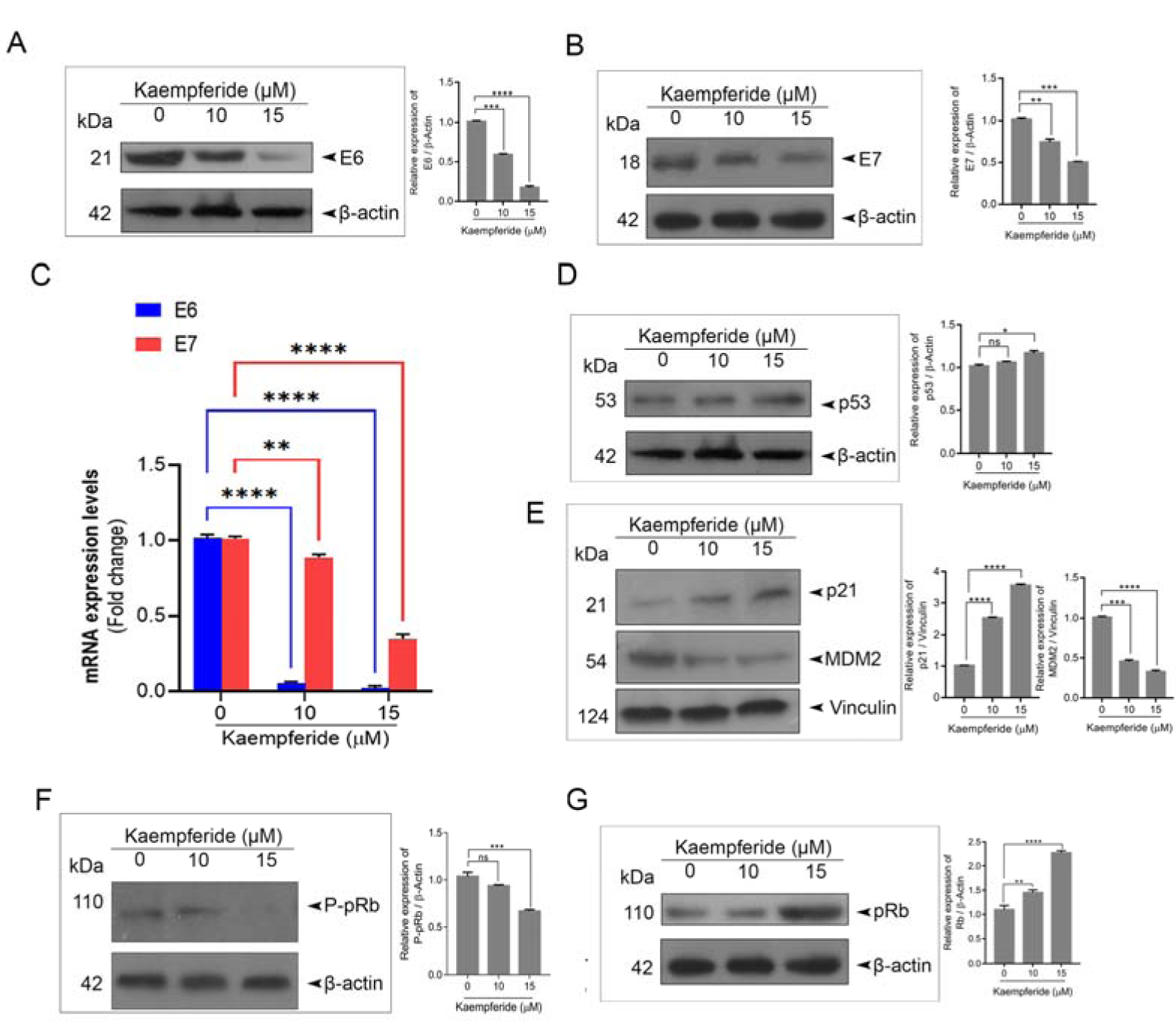
Kaempferide inhibits HPV oncoproteins E6 and E7 leading to p53 and pRb-mediated apoptosis: **(A-B)** Western blot showing that kaempferide down-regulates the oncoproteins, E6 and E7. **(C)** qPCR analysis confirms the efficacy of kaempferide to down-regulate E6 and E7. **(D-E)** Western blot analysis demonstrating that kaempferide induces p53 and p21, while down-regulating MDM2 expression. [Data are representative of three independent experiments (Mean±SEM) and P-values are calculated using two way ANOVA****P ≤0.0001,***P ≤0.001,**P ≤0.01,*P ≤0.1and ns ≥ 0.05] **(F-G)** Western blot revealing the efficacy of kaempferide to up-regulate pRb and degrade phospho-pRb. [Data are representative of three independent experiments (Mean±SEM) and P-values are calculated using one way ANOVA****P ≤0.0001,***P ≤0.001,**P ≤0.01,*P ≤0.1and ns ≥ 0.05].

### 3.2 Kaempferide inhibits development of cervical tumor xenografts in NOD-SCID mice by inducing apoptosis through down-regulation of E6 and E7

To evaluate the efficacy of kaempferide against cervical cancer, *in vivo*, a cervical cancer xenograft model developed in NOD-SCID (NOD.CB17-prkdcscid/J) mice bearing Hela-Luc+ xenografts was used. The schematic of the human cervical cancer development and the drug treatment regimen in the NOD-SCID murine model have been illustrated in **Figure 3A** and explained in the methodology. Following the drug treatment, the animals were euthanized and tumors were procured for further analysis. The tumor volume and luc activity were drastically reduced in kaempferide-treated mice, compared to that of the control. While the mean tumor volume of the control group was 1126mm^3^, it was 289mm^3^ in the kaempferide-treated group at the time of sacrifice **(Figure 3B-D)**. Histopathological analysis of the excised tumor samples from the control group revealed tumor cells with active mitosis while tissues of kaempferide-treated mice displayed massive apoptosis, authenticating that kaempferide induces apoptotic mode of cell death in cervical cancer tissues **(Figure 3E)**. Furthermore, immunohistochemical analysis for PARP and TUNEL staining revealed a drastic increase in the number of apoptotic cells in the kaempferide-treated xenografts in comparison to the control (**Figure 3F-G)**.

**Figure 3:**
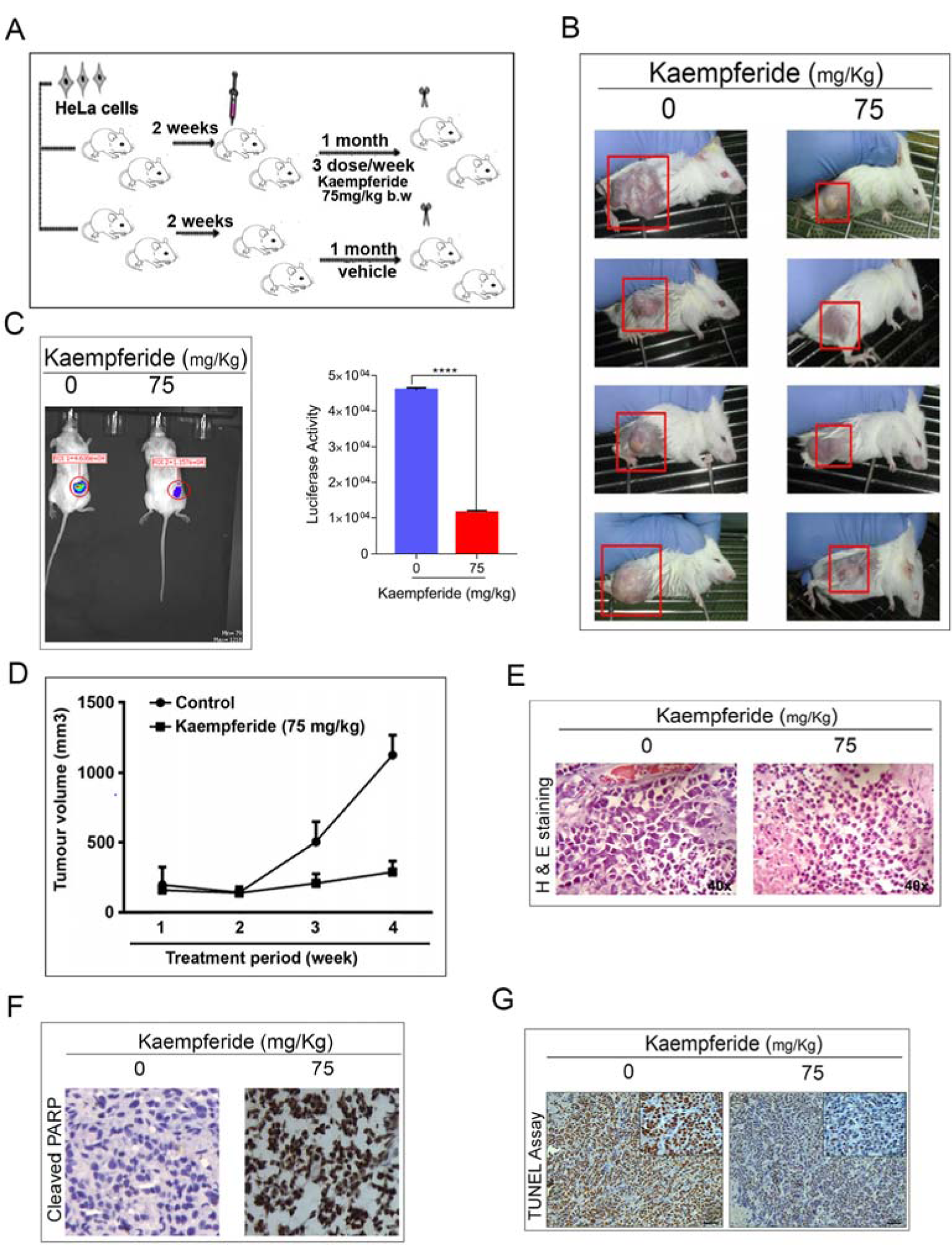
Kaempferide inhibits the growth of cervical cancer xenografts inducing apoptotic mode of cell death. **(A)** A schematic depicting the human cervical cancer xenograft model in NOD-SCID mice using HeLa cells and the drug treatment regimen. **(B)** Representative images of control and kaempferide-treated mice bearing HeLa xenografts. **(C)** Representative image showing the difference in Luc intensity in control and kaempferide-treated mice. **(D)** A graphical representation of tumor volumes of indicated treatment groups. **(E)** Histopathological staining of xenograft-derived tumors from control and treatment groups. **(F)** Immunohistochemical analysis of apoptosis marker, cleaved PARP in tumor sections. **(G)** Increase in TUNEL-positive cells following kaempferide-treatment confirms apoptotic mode of cell death.

A significant down-regulation of E6 and E7 and subsequent up-regulation of p53, p21, and a significant down-regulation of MDM2 was noted in xenograft tissues of kaempferide-treated mice **(Figure 4A-D)**. Degradation of phospho-pRb with simultaneous up-regulation of pRb was also noted in the kaempferide-treated cervical cancer xenografts, attesting the in vitro results displayed in Figure 1 **(Figure 4E, F)**. As observed in the *in vitro* studies, immunohistochemical analysis of the kaempferide-treated tumor tissues for the expression status of the cell proliferation marker, PCNA and the HPV oncoproteins, E6 and E7 displayed a significant down-regulation. While E6 induces cervical cancer progression through degradation of p53, E7 does the same by suppression of pRb. We noted a substantial up-regulation of the apoptosis markers, p53 and p21 and down-regulation of the oncoprotein, MDM2. Corresponding to the down-regulation of E7, an up-regulation of pRb, the downstream target of E7 was also noted, corroborating the therapeutic potential of kaempferide against cervical cancer **(Figure 4G)**.

**Figure 4:**
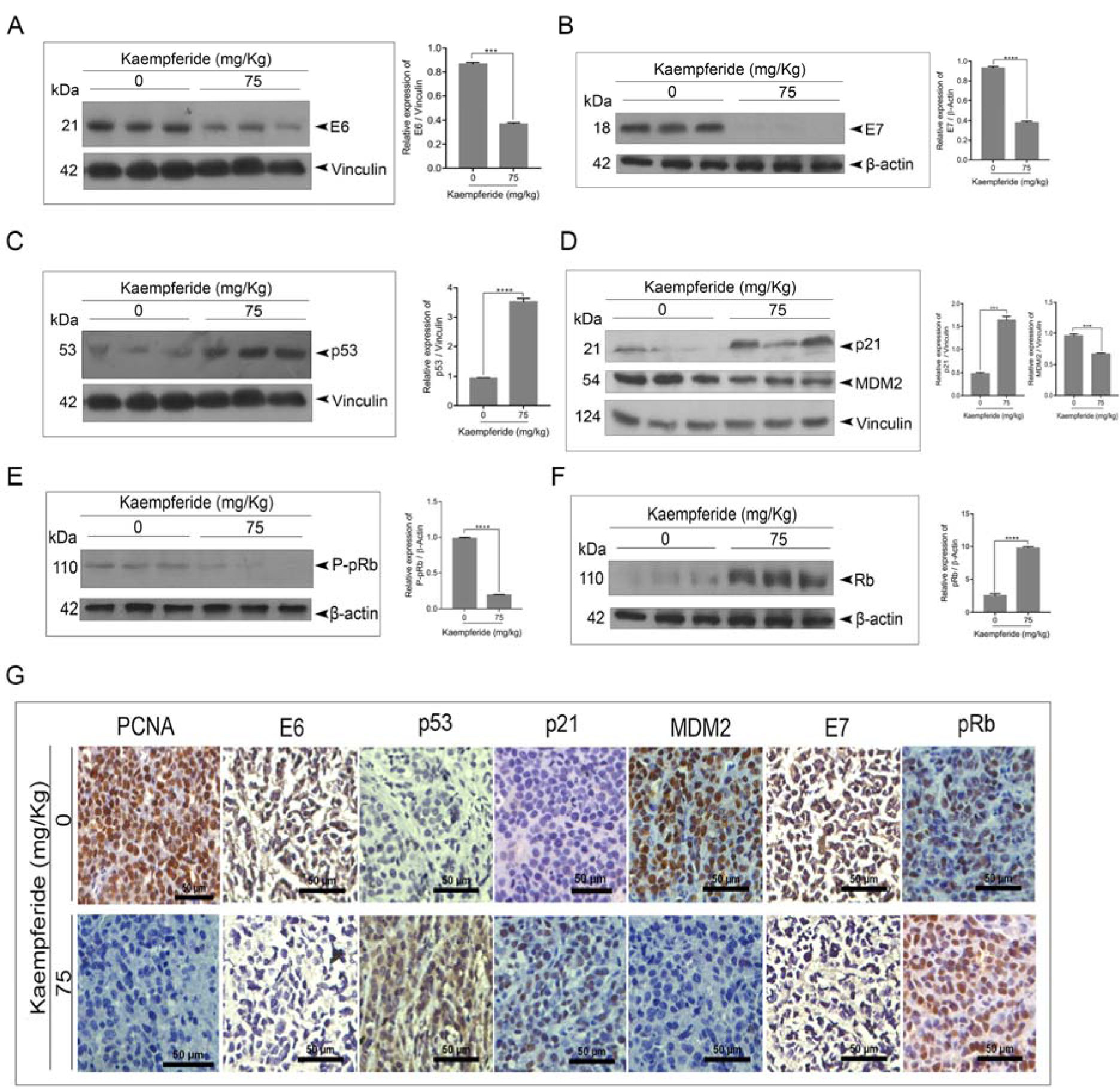
Kaempferide inhibits the growth of cervical cancer xenografts by down-regulating the HPV oncoproteins, leading to apoptotic mode of cell death. **(A-B)** Western blot showing the down-regulation of HPV oncoproteins, E6 and E7 in response to kaempferide-treatment in HeLa xenografts. (**C-D**) Western blot analysis reveals that kaempferide up-regulates tumor suppressor proteins, p53 and p21 and down-regulates the oncoprotein, MDM2. (**E-F**) Western blot demonstrating that kaempferide degrades phospho-pRb leading to the over-expression of pRb. (**G**) Immunohistochemical analysis illustrating that kaempferide inhibits the nuclear proliferation marker PCNA, and down-regulates the HPV oncoproteins, E6 and E7 leading to the up-regulation of tumor suppressor proteins, p53, p21 and pRb and degradation of the oncoprotein MDM2, in tumor sections. [Data are representative of three independent experiments (Mean±SEM) and P-values are calculated using one way ANOVA****P ≤0.0001, ***P ≤0.001, **P ≤0.01, *P ≤0.1and ns ≥ 0.05].

## 4. Discussion

According to WHO statistics, despite the invention of preventive vaccines and cytology-based cervical cancer prevention programs, cervical cancer is still the foremost reason for cancer-related death among women in developing countries. Based on a worldwide meta-analysis study, approximately 291 million women have been estimated to have HPV infection [22]. It has been suggested that up to 80% of sexually active women will acquire an HPV infection at some point during their lifetime. Although approval of the Merck vaccine, Gardasil has proceeded rapidly in many countries, it is still unaffordable to patients from developing countries, who are the real victims of cervical cancer. Though several chemotherapeutic drugs have been developed for the treatment of cervical cancer, most of them have either severe side effects or cost unaffordable to the patients, who are most often having a poor socio-economic status. Since many of the currently available chemotherapeutic agents are phytochemicals, plants can be a potential source for exploring new drugs. Being a group bioprospecting for anticancer natural products, we have reported some potent anticancer molecules [23], [24]. We had previously reported a very effective chemotherapeutic combinatorial regimen of paclitaxel and the phytochemical, curcumin, with remarkably lesser cost as well as side effects, for the treatment of cervical cancer [18, 25, 26]. Ours is the first study reporting the anticancer potential and pharmacological safety of kaempferide [17]. However, other groups have reported the anti-proliferative activity of kaempferide triglycoside and kaempferide mannich bases [27, 28]. The present study is the first *in vivo* validation of the anti-cancer efficacy of kaempferide, using a human cervical cancer xenograft model in NOD-SCID mice. Antitumor effect of kaempferide was analyzed on ectopically grown human cervical cancer in the flank region of NOD-SCID mice. The reduction in tumor volume and IHC analysis of PCNA in tumor cryosections of kaempferide-treated mice compared to that of untreated controls demonstrate the efficacy of this compound against cervical cancer. The enhancement of apoptosis in cervical cancer xenograft tissues treated with kaempferide as evidenced by the high expression of cleaved PARP in immunohistochemistry and high positivity in TUNEL staining, were in concordance with our previous *in vitro* results [17].

In the present study, we have investigated the ability of kaempferide to down-regulate the HPV oncoproteins, E6 and E7. E6 is known to induce cervical cancer progression through degradation of p53, while E7 does the same by suppression of pRb and/or by activating the cellular oncoprotein, MDM2, which promotes the ubiquitination and proteasomal degradation of p53, by acting as a ubiquitin ligase [29]. Studies indicate that cells expressing E6, whether alone or in combination with E7 have decreased levels of p53 due to the ability of E6 to facilitate its degradation [20]. As p53 in HeLa is reported as minimal due to E6 over-expression, we examined the effect of kaempferide on E6 and E7 expression status. Interestingly, kaempferide induced strong E6 down-regulation, which may be the reason for the subsequent up-regulation of p53 and its down-stream signaling molecule, p21. Mdm2 is a major mechanism utilized by cancer cells to escape p53 surveillance. This oncoprotein has been shown to complex with p53 and sterically inhibit p53 transcriptional activity. Mdm2 also inhibits p53 activities by promoting p53 nuclear export. Mdm2-p53 signaling is regulated during the DNA damage response by posttranslational modification of p53 [30]. Both *in vitro* and *in vivo* results revealed a significant down-regulation of MDM2 levels in response to kaempferide. The compound also induced a significant down-regulation of E7 expression, which possibly led to the degradation of phospho-pRb and subsequent over-expression of pRb. pRb acts as a transcriptional co-repressor, which inhibits expression of multiple genes necessary for cell cycle progression leading to induction of cell death in cervical cancer cells. By inactivating pRb via promoting its degradation through phosphorylation of pRb, E7 circumvents the tumor suppressive function of pRb, subsequently promoting uncontrolled cell division [31, 32]. Hence kaempferide, through inhibition of E7, degrades the phospho-pRb to retrieve the pRb function, which leads to induction of apoptosis in cervical cancer cells.

Taken together, our study reveals the antioncogenic potential of kaempferide, which manifests its efficacy through p53 and pRb, by down-regulating the HPV proteins, E6 and E7. E6 and E7 are the key oncoproteins regulating the process of HPV-mediated cervical tumorigenesis by establishing all the major hallmarks of cancer [33]. Hence, they are the most effective targets for cervical cancer therapeutics, and effective management of their expression status can ensure the eradication of all cervical cancer cells. It is in this scenario our findings on kaempferide should gain attention. This is the first report, which demonstrates kaempferide as an inhibitor of HPV oncoproteins and hence as a promising drug molecule against cervical cancer.

In conclusion, the present study demonstrates with mechanism based evidence, the effectiveness of kaempferide against cervical cancer. Since it verifies our previous *in vitro* results on the anti-cancer efficacy of kaempferide against cervical cancer through *in vivo* cervical xenograft model, this study can be considered as the first pre-clinical intervention on this compound as a candidate drug against human cervical cancer.

## Author Contributions

Study concept and design: RJA. Acquisition, analysis, or interpretation of the data: LRN, VVA, SUA, TPR, NHH, CKK, AKTT, MS, RSL. Drafting of the paper: VVA, TPR, CKK, NHH. Figures: SUA. Critical review of the manuscript: RJA. All authors have read and agreed to the current version of the manuscript.

## Funding

This research was funded by Kerala State Council for Science, Technology and Engineering, Government of Kerala (Funding Grant No.008/SRSLS/2014/CSTE).

## Institutional Review Board Statement

The animal study protocol was approved by the Institutional Review Board of Rajiv Gandhi Centre for Biotechnology (IAEC/721/RUBY/2018 approved on 9-12-2019).

## Data Availability Statement

Raw data compliant with the institutional confidentiality policies are available upon request. Data requests should be sent to the corresponding author. Data is contained within the article.

## Acknowledgments

We acknowledge the immense help provided by the RGCB central instrumentation facilities for the successful completion of the experiments.

## Conflicts of Interest

The authors declare no competing interest. The funders had no role in the design of the study; in the collection, analyses, or interpretation of data; in the writing of the manuscript; or in the decision to publish the results.

## References

[1] M. Schiffman, N. Wentzensen, S. Wacholder, W. Kinney, J.C. Gage, P.E. Castle, Human papillomavirus testing in the prevention of cervical cancer, Journal of the National cancer institute 103(5) (2011) 368–383.

[2] B.A. Werness, A.J. Levine, P.M. Howley, Association of human papillomavirus types 16 and 18 E6 proteins with p53, Science 248(4951) (1990) 76-79.

[3] Z. Hu, W. Ding, D. Zhu, L. Yu, X. Jiang, X. Wang, C. Zhang, L. Wang, T. Ji, D. Liu, D. He, X. Xia, T. Zhu, J. Wei, P. Wu, C. Wang, L. Xi, Q. Gao, G. Chen, R. Liu, K. Li, S. Li, S. Wang, J. Zhou, D. Ma, H. Wang, TALEN-mediated targeting of HPV oncogenes ameliorates HPV-related cervical malignancy, The Journal of clinical investigation 125(1) (2015) 425–36.

[4] Z. Hu, L. Yu, D. Zhu, W. Ding, X. Wang, C. Zhang, L. Wang, X. Jiang, H. Shen, D. He, K. Li, L. Xi, D. Ma, H. Wang, Disruption of HPV16-E7 by CRISPR/Cas system induces apoptosis and growth inhibition in HPV16 positive human cervical cancer cells, BioMed research international 2014 (2014) 612823.

[5] C.H. Yuan, M. Filippova, S.S. Tungteakkhun, P.J. Duerksen-Hughes, J.L. Krstenansky, Small molecule inhibitors of the HPV16-E6 interaction with caspase 8, Bioorganic & medicinal chemistry letters 22(5) (2012) 2125–9.

[6] L. Wang, H. Guo, L. Yang, L. Dong, C. Lin, J. Zhang, P. Lin, X. Wang, Morusin inhibits human cervical cancer stem cell growth and migration through attenuation of NF-κB activity and apoptosis induction, Molecular and cellular biochemistry 379(1-2) (2013) 7–18.

[7] J.H. Jung, J.O. Lee, J.H. Kim, S.K. Lee, G.Y. You, S.H. Park, J.M. Park, E.K. Kim, P.G. Suh, J.K. An, H.S. Kim, Quercetin suppresses HeLa cell viability via AMPK-induced HSP70 and EGFR down-regulation, Journal of cellular physiology 223(2) (2010) 408–14.

[8] K.S. Chung, J.H. Choi, N.I. Back, M.S. Choi, E.K. Kang, H.G. Chung, T.S. Jeong, K.T. Lee, Eupafolin, a flavonoid isolated from Artemisia princeps, induced apoptosis in human cervical adenocarcinoma HeLa cells, Molecular nutrition & food research 54(9) (2010) 1318–28.

[9] A.K. Jha, M. Nikbakht, G. Parashar, A. Shrivastava, N. Capalash, J. Kaur, Reversal of hypermethylation and reactivation of the RARβ2 gene by natural compounds in cervical cancer cell lines, Folia biologica 56(5) (2010) 195–200.

[10] A.J. Alonso-Castro, E. Ortiz-Sánchez, A. García-Regalado, G. Ruiz, J.M. Núñez-Martínez, I. González-Sánchez, V. Quintanar-Jurado, E. Morales-Sánchez, F. Dominguez, G. López-Toledo, M.A. Cerbón, A. García-Carrancá, Kaempferitrin induces apoptosis via intrinsic pathway in HeLa cells and exerts antitumor effects, Journal of ethnopharmacology 145(2) (2013) 476–89.

[11] K. Bishayee, S. Ghosh, A. Mukherjee, R. Sadhukhan, J. Mondal, A.R. Khuda-Bukhsh, Quercetin induces cytochrome-c release and ROS accumulation to promote apoptosis and arrest the cell cycle in G2/M, in cervical carcinoma: signal cascade and drug-DNA interaction, Cell proliferation 46(2) (2013) 153–63.

[12] X. Yuan, B. Zhang, N. Chen, X.Y. Chen, L.L. Liu, Q.S. Zheng, Z.P. Wang, Isoliquiritigenin treatment induces apoptosis by increasing intracellular ROS levels in HeLa cells, Journal of Asian natural products research 14(8) (2012) 789–98.

[13] M. Singh, S. Tyagi, K. Bhui, S. Prasad, Y. Shukla, Regulation of cell growth through cell cycle arrest and apoptosis in HPV 16 positive human cervical cancer cells by tea polyphenols, Investigational new drugs 28 (2010) 216–224.

[14] H.G. Lee, K.A. Yu, W.K. Oh, T.W. Baeg, H.C. Oh, J.S. Ahn, W.C. Jang, J.W. Kim, J.S. Lim, Y.K. Choe, D.Y. Yoon, Inhibitory effect of jaceosidin isolated from Artemisiaargyi on the function of E6 and E7 oncoproteins of HPV 16, Journal of ethnopharmacology 98(3) (2005) 339–43.

[15] Y. Qiao, J. Cao, L. Xie, X. Shi, Cell growth inhibition and gene expression regulation by (-)-epigallocatechin-3-gallate in human cervical cancer cells, Archives of pharmacal research 32(9) (2009) 1309–15.

[16] S. Ham, K.H. Kim, T.H. Kwon, Y. Bak, D.H. Lee, Y.S. Song, S.H. Park, Y.S. Park, M.S. Kim, J.W. Kang, J.T. Hong, D.Y. Yoon, Luteolin induces intrinsic apoptosis via inhibition of E6/E7 oncogenes and activation of extrinsic and intrinsic signaling pathways in HPV-18-associated cells, Oncology reports 31(6) (2014) 2683–91.

[17] L.R. Nath, J.N. Gorantla, S.M. Joseph, J. Antony, S. Thankachan, D.B. Menon, S. Sankar, R.S. Lankalapalli, R.J.J.R.a. Anto, Kaempferide, the most active among the four flavonoids isolated and characterized from Chromolaena odorata, induces apoptosis in cervical cancer cells while being pharmacologically safe, 5(122) (2015) 100912–100922.

[18] C.N. Sreekanth, S.V. Bava, E. Sreekumar, R.J. Anto, Molecular evidences for the chemosensitizing efficacy of liposomal curcumin in paclitaxel chemotherapy in mouse models of cervical cancer, Oncogene 30(28) (2011) 3139–52.

[19] T. Kobayashi, T. Makino, K. Yamashita, T. Saito, K. Tanaka, T. Takahashi, Y. Kurokawa, M. Yamasaki, K. Nakajima, E. Morii, APR-246 induces apoptosis and enhances chemo-sensitivity via activation of ROS and TAp73-Noxa signal in oesophageal squamous cell cancer with TP53 missense mutation, British Journal of Cancer 125(11) (2021) 1523–1532.

[20] J.T. Thomas, L.A. Laimins, Human papillomavirus oncoproteins E6 and E7 independently abrogate the mitotic spindle checkpoint, Journal of virology 72(2) (1998) 1131–1137.

[21] E. Berezutskaya, S. Bagchi, The human papillomavirus E7 oncoprotein functionally interacts with the S4 subunit of the 26 S proteasome, Journal of Biological Chemistry 272(48) (1997) 30135–30140.

[22] S.V. Bava, A.K.T. Thulasidasan, C.N. Sreekanth, R.J. Anto, Cervical cancer: A comprehensive approach towards extermination, Annals of medicine 48(3) (2016) 149–161.

[23] A. Shabna, J. Antony, V. Vijayakurup, M. Saikia, V.B. Liju, A.P. Retnakumari, N.A. Amrutha, V.V. Alex, M. Swetha, S.U. Aiswarya, Pharmacological attenuation of melanoma by tryptanthrin pertains to the suppression of MITF-M through MEK/ERK signaling axis, Cellular and Molecular Life Sciences 79(9) (2022) 478.

[24] L.R. Nath, J.N. Gorantla, A.K.T. Thulasidasan, V. Vijayakurup, S. Shah, S. Anwer, S.M. Joseph, J. Antony, K.S. Veena, S. Sundaram, Evaluation of uttroside B, a saponin from Solanum nigrum Linn, as a promising chemotherapeutic agent against hepatocellular carcinoma, Scientific reports 6(1) (2016) 36318.

[25] S.V. Bava, V.T. Puliappadamba, A. Deepti, A. Nair, D. Karunagaran, R.J. Anto, Sensitization of taxol-induced apoptosis by curcumin involves down-regulation of nuclear factor-κB and the serine/threonine kinase Akt and is independent of tubulin polymerization, Journal of Biological Chemistry 280(8) (2005) 6301–6308.

[26] A.K.T. Thulasidasan, A.P. Retnakumari, M. Shankar, V. Vijayakurup, S. Anwar, S. Thankachan, K.S. Pillai, J.J. Pillai, C.D. Nandan, V.V. Alex, Folic acid conjugation improves the bioavailability and chemosensitizing efficacy of curcumin-encapsulated PLGA-PEG nanoparticles towards paclitaxel chemotherapy, Oncotarget 8(64) (2017) 107374.

[27] V. Martineti, I. Tognarini, C. Azzari, S.C. Sala, F. Clematis, M. Dolci, V. Lanzotti, F. Tonelli, M.L. Brandi, P. Curir, Inhibition of in vitro growth and arrest in the G0/G1 phase of HCT8 line human colon cancer cells by kaempferide triglycoside from Dianthus caryophyllus, Phytotherapy Research 24(9) (2010) 1302–1308.

[28] V.-S. Nguyen, L. Shi, F.-Q. Luan, Q.-A. Wang, Synthesis of kaempferide Mannich base derivatives and their antiproliferative activity on three human cancer cell lines, Acta Biochimica Polonica 62(3) (2015).

[29] M. Traidej, L. Chen, D. Yu, S. Agrawal, J. Chen, The roles of E6-AP and MDM2 in p53 regulation in human papillomavirus-positive cervical cancer cells, Antisense and Nucleic Acid Drug Development 10(1) (2000) 17–27.

[30] S. Hietanen, S. Lain, E. Krausz, C. Blattner, D.P. Lane, Activation of p53 in cervical carcinoma cells by small molecules, Proceedings of the National Academy of Sciences 97(15) (2000) 8501–8506.

[31] R.J. Bourgo, W.A. Braden, S.I. Wells, E.S. Knudsen, Activation of the retinoblastoma tumor suppressor mediates cell cycle inhibition and cell death in specific cervical cancer cell lines, Molecular Carcinogenesis: Published in cooperation with the University of Texas MD Anderson Cancer Center 48(1) (2009) 45-55.

[32] M.V. Frolov, N.J. Dyson, Molecular mechanisms of E2F-dependent activation and pRB-mediated repression, Journal of cell science 117(11) (2004) 2173–2181.

[33] A. Pal, R. Kundu, Human papillomavirus E6 and E7: the cervical cancer hallmarks and targets for therapy, Frontiers in microbiology 10 (2020) 3116.

